# Enhancing Precision in Bioprinting Utilizing Fuzzy Systems

**DOI:** 10.1101/2021.07.10.451921

**Authors:** Ashkan Sedigh, Dayna DiPiero, Kristy M. Shine, Ryan E. Tomlinson

## Abstract

Bioprinting facilitates the generation of complex, three-dimensional (3D), cell-based constructs for a variety of applications. Although multiple bioprinting technologies have been developed, extrusion-based systems have become the dominant technology due to the diversity of substrate materials (bioinks) that can be accommodated, either individually or in combination. However, each bioink has unique material properties and extrusion characteristics that limit bioprinting precision, particularly when generating constructs from different bioinks. Here, we aimed to achieve high precision (i.e. repeatability) across samples by generating bioink-specific printing parameters using a systematic approach. We hypothesized that a fuzzy system could be used as a “black box” method to tackle the inherent vagueness and imprecision in 3D bioprinting data and uncover the optimal printing parameters for a specific bioink that would result in high accuracy and precision. Our fuzzy model was used to approximate and quantify the precision and ease of printability for two common bioinks - type I collagen and Pluronic F127, with or without dilution in αMEM culture media. The model consisted of three inputs (pressure, speed, and dilution percent of bioink) and a single output (layer width). Using this system, we introduce the Bioink Precision Index (BPI), a metric that can be used to quantify and compare the precision of any bioink. Here, we show that printing with parameters optimized using BPI increases the precision for collagen (+15%) and Pluronic F127 (+29%) as compared to the manufacturer’s recommended printing parameters.

## 1. Introduction

Bioprinting is a popular technique used in a variety of research areas, such as tissue engineering and drug delivery, and involves depositing biological material (bioinks) in a layer-by-layer fashion to produce a three-dimensional (3D) cell-laden construct [1]. The generation of bioprinted 3D constructs has opened the door to many novel applications, including visualization of cell-cell interactions in the natural 3D microenvironment for cancer research [2], production of 3D tissue for implantation [3], and in vivo models for drug discovery [4]. Bioprinted constructs are designed and generated from a computer-aided design (CAD) [5], and can thus be reproduced quickly from a vast and expanding library of bioinks [6], [7]. As a result, methods that increase precision and facilitate reproducibility are particularly desirable [8], [9].

Extrusion-based bioprinting (EBB) is the most utilized bioprinting technique due to its compatibility with a large spectrum of bioinks, affordability, and ease of use [10]. In this method, bioinks are extruded from a small nozzle in a layer-by-layer manner using pressure generated from either a pneumatic, screw, or piston system [10], [11]. As a result, the bioprinting parameters, such as nozzle pressure, nozzle gauge, printing speed, material temperature, and crosslinking status, affect the final construct [8]. Moreover, these parameters must be optimized for each bioink in order to generate a final construct that has the desired dimensions and appropriate material properties.

Type I collagen and Pluronic F-127 bioinks (also known as Poloxamer 407) are two widely used bioinks in biomedical research. Collagen bioink is a biocompatible, protein-based hydrogel bioink that is commonly found in the extracellular matrix in a variety of tissues [12]. Importantly, collagen bioink must be extruded below 4 °C, since it irreversibly gels at higher temperatures and cannot be brought back to the liquid state [10], [12]. In contrast, Pluronic F-127 is a synthetic hydrogel consisting of a co-polymer tri-block structure with hydrophilic-hydrophobic-hydrophilic sequences [13]. Similar to collagen bioink, Pluronic F-127 is aqueous at 4 °C, but gels at room temperature. However, Pluronic F-127 can be returned to a liquid state by cooling the bioprinted construct, leading to its common use as a sacrificial support material for forming hollow structures or delivering cells [12]–[15].

Few systematic approaches have been developed to approximate optimized bioprinting parameters [16]. The disadvantage of using mathematical modelling approaches is a lack of generalization to more than a specific number of inputs. Moreover, the linearity of the modeling may lead to imprecision and inaccuracy in output approximation for new bioinks formulations [17]. To overcome these shortcomings, we hypothesized that bioprinting parameter optimization could be robustly executed by implementing a fuzzy logic system, in which rules are defined to assign the input and outputs continuous values, rather than the discrete values generally utilized [18].

To test this hypothesis, we developed a fuzzy system to optimize bioprinting parameters for collagen and Pluronic F-127 bioinks with or without dilution in αMEM culture media. Our fuzzy system consists of inputs of nozzle pressure, printing speed, and dilution percentage of bioink with a single output of layer width. The results from our study suggest that this approach is useful for optimizing printing parameters and will improve reproducibility across diverse bioinks as well as provide an objective characterization of bioprinting precision for newly formulated bioinks.

## 2. Methods

### 2.1. Bioprinting

Pluronic F-127 and type I collagen bioinks (Allevi Collagen Lifeink® 200, *USA*) were prepared for analysis by diluting with alpha Minimum Essential Medium (αMEM) to contain 0, 20, or 40% of media, to mimic potential applications in which cells would be seeded into each bioink. The dilutions were performed by mixing between two syringes connected with a female-to-female Luer Lock coupler. To confirm the material was homogenously mixed, the Pluronic F-127 used for this study was colored using 600 uL of commercially available food dye.

Each bioink was used to print a 1×1 cm square shape with a layer width and height of 200 microns (Fig. 1A). For each bioink, we used a 0.25 inch straight 25 gauge nozzle. Our initial experimentation was performed with undiluted bioinks (dilution held constant at 0%). For the collagen bioink, we tested three extrusion pressures (15, 20, 25 psi) and five printing speeds (8, 10, 12, 14, 15 mm/s). For the Pluronic F-127, we tested four extrusion pressures (60, 70, 80, 90 psi) and four printing speeds (12, 13, 14, 15 mm/s). Next, to determine the effect of dilution, we used a constant printing speed (15 mm/s). For the collagen bioink, we tested 20% and 40% diluted collagen at three extrusion pressures (8, 10, 12 psi). For the Pluronic F-127, we tested three extrusion pressures at 20% dilution (40, 45, 50 psi) and 40% dilution (20, 25, 35 psi). All studies were performed using three independent experiments. Since collagen needs to be printed at 4°C or below, an ice pack was secured to the extruder throughout the print (Fig. 1B). For all experiments, we used an Allevi 2 bioprinter (*Allevi, USA*) located inside a sterile biosafety cabinet.

**Figure 1.**
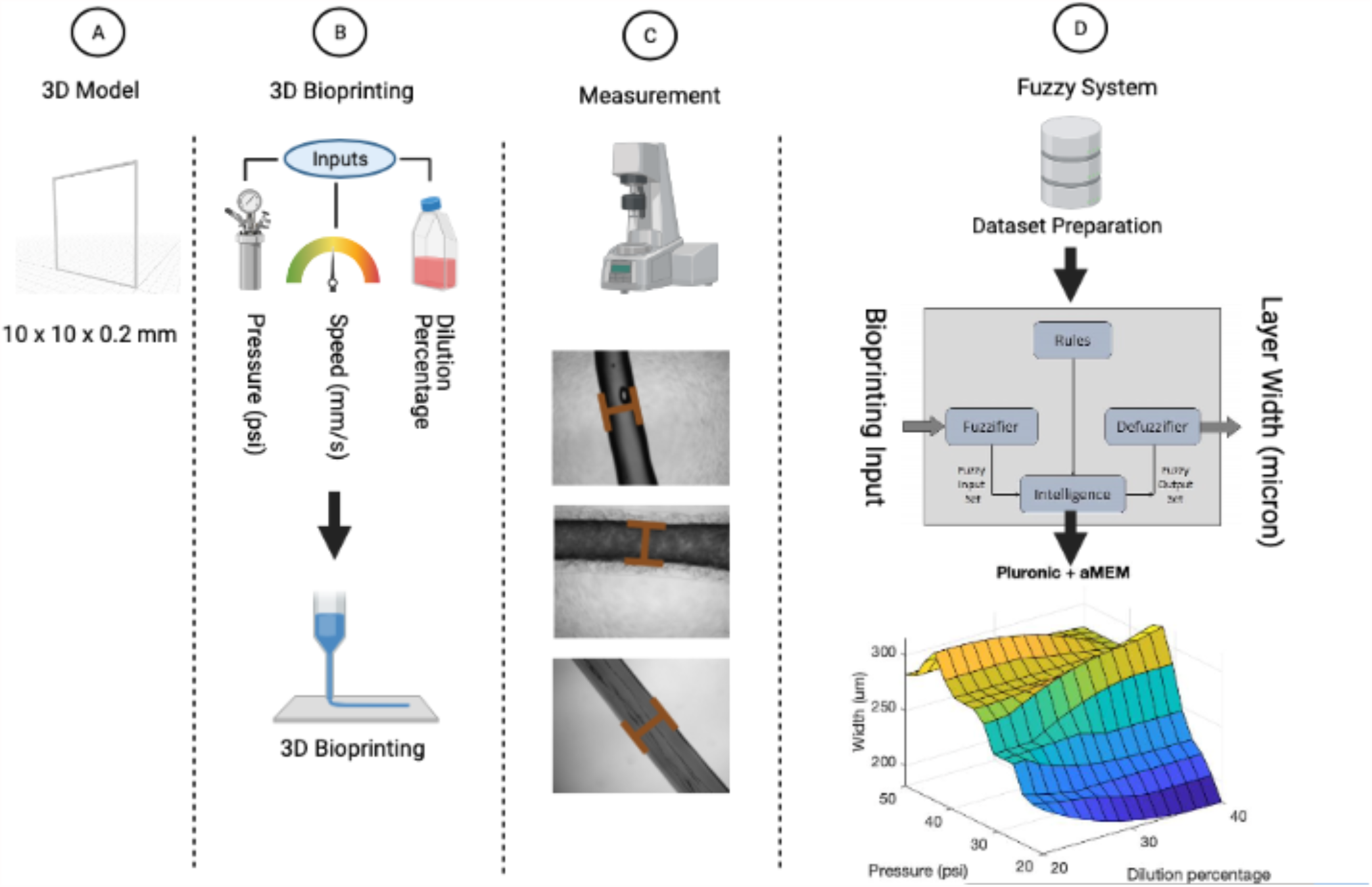
General workflow of the proposed study. **(A)** is the first step that the 3D model is designed and with three different input parameters the sample is printed by an extrusion based bioprinter (B). In step (C) data is measured, processed and fed into the fuzzy system rules, then the final 3D surface is generated to calculate the BP for the preferred points (D)

### 2.2. Imaging and Analysis

First, each sample was imaged using brightfield microscopy at 4x objective size using a Nikon E800 microscope (Fig. 1C). Next, images were analyzed in FIJI [19] by quantifying the width of the layer at the midpoint of each side of the square. The values from four independent trials were averaged to obtain the final value for each sample. These measurements were then utilized as the outputs for the inference engine in the fuzzy system (Fig. 1D).

### 2.3. Implementing Fuzzy System

We implemented a fuzzy system to utilize our experimental data to optimize bioprinting parameters, as previously described [20]. Briefly, we defined our crisp input values to be pressure, speed, and dilution percentage with layer width as a single output value. The process of converting these crisp inputs to fuzzy values is known as fuzzification. Here, we used Gaussian membership functions, which are popular because of their smoothness, concise notation, and similarity to a variety of biological processes [21]. Next, mapping from a given fuzzy input to a fuzzy output was performed using a Mamdani fuzzy inference system (Fuzzy Logic Toolbox, Matlab R2020a). In this process, we imported a series of *If-Then* rules (experimental results) to the inference engine (Tables 1-4). Finally, a single value for the output (layer width) can be generated as an aggregate of fuzzy values in a process known as defuzzification. A schematic of our fuzzy system approach is illustrated in Figure 2. We note that our fuzzy system is based on the following assumptions: (i) the output material is incompressible, (ii) the pressure drop during extrusion is negligible, (iii) the flow is steady and laminar.

**Table 1.** Type-1 Fuzzy system rules for Pluronic experiment

| Rule | Pressure (psi)<br>SD: 4.24 | Speed (mm/s)<br>SD: 0.42 | Output layer width (micron)<br>SD:16.99 |
| --- | --- | --- | --- |
| 1 | 60 | 12 | 180 |
| 2 | 60 | 13 | 220 |
| 3 | 60 | 14 | 180 |
| 4 | 60 | 15 | 180 |
| 5 | 70 | 12 | 300 |
| 6 | 70 | 13 | 260 |
| 7 | 70 | 14 | 260 |
| 8 | 70 | 15 | 220 |
| 9 | 80 | 12 | 260 |
| 10 | 80 | 13 | 300 |
| 11 | 80 | 14 | 300 |
| 12 | 80 | 15 | 260 |
| 13 | 90 | 12 | 420 |
| 14 | 90 | 13 | 340 |
| 15 | 90 | 14 | 340 |
| 16 | 90 | 15 | 340 |

**Table 2.** Type-1 Fuzzy system rules for Collagen experiment

| Rule | Pressure (psi)<br>SD: 2.12 | Speed (mm/s)<br>SD: 0.74 | Output layer width (micron)<br>SD:42.4 |
| --- | --- | --- | --- |
| 1 | 15 | 8 | 350 |
| 2 | 15 | 10 | 250 |
| 3 | 15 | 12 | 250 |
| 4 | 15 | 14 | 350 |
| 5 | 15 | 15 | 250 |
| 6 | 20 | 8 | 850 |
| 7 | 20 | 10 | 650 |
| 8 | 20 | 12 | 650 |
| 9 | 20 | 14 | 550 |
| 10 | 20 | 15 | 550 |
| 11 | 25 | 8 | 950 |
| 12 | 25 | 10 | 950 |
| 13 | 25 | 12 | 950 |
| 14 | 25 | 14 | 850 |
| 15 | 25 | 15 | 850 |

**Table 3.** Type-1 Fuzzy system rules for Diluted Collagen with aMEM experiment

| Rule | Dilution Percentage<br>SD:8.43 | Pressure (psi)<br>SD: 0.84 | Output layer width (micron)<br>SD: 21.24 |
| --- | --- | --- | --- |
| 1 | 20 | 8 | 250 |
| 2 | 20 | 10 | 200 |
| 3 | 20 | 12 | 300 |
| 4 | 40 | 8 | 200 |
| 5 | 40 | 10 | 350 |
| 6 | 40 | 12 | 400 |

**Table 4.**
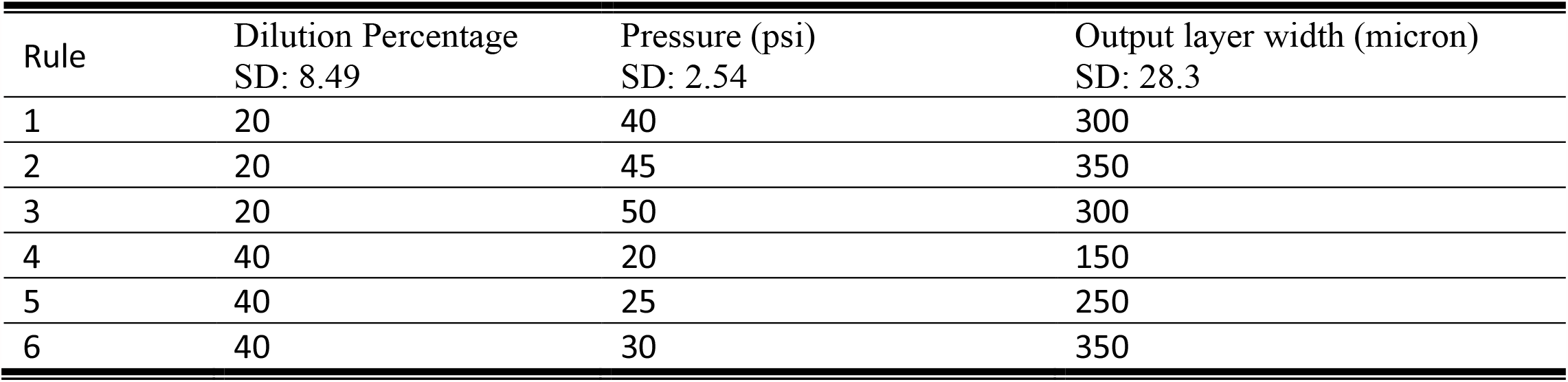
Type-1 Fuzzy system rules for the Diluted Pluronic with aMEM experiment

| Rule | Dilution Percentage<br>SD: 8.49 | Pressure (psi)<br>SD: 2.54 | Output layer width (micron)<br>SD: 28.3 |
| --- | --- | --- | --- |
| 1 | 20 | 40 | 300 |
| 2 | 20 | 45 | 350 |
| 3 | 20 | 50 | 300 |
| 4 | 40 | 20 | 150 |
| 5 | 40 | 25 | 250 |
| 6 | 40 | 30 | 350 |

**Figure 2.**
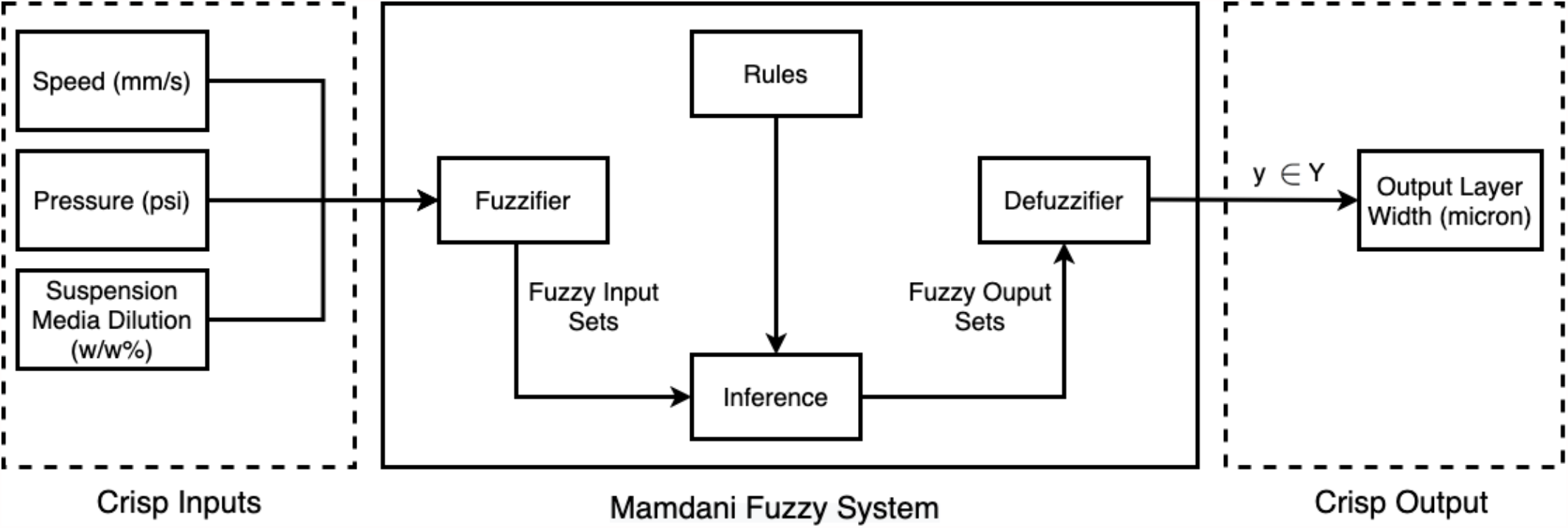
Type-1 Fuzzy Logic Algorithm and Study Design. General overview and different features of the Type-1 Fuzzy system including fuzzification, rules, inference engine, and defuzzification. It includes three inputs (speed, pressure, and Suspension Media Dilution Factor) and one output (layer width).

## 3. Results

### 3.1. Imprecision in undiluted bioink is primarily due to nozzle pressure, not printing speed

First, we utilized our fuzzy system to visualize the layer width (output) as a function of the inputs, which were printing speed (mm/s) and nozzle pressure (psi) in collagen bioink and Pluronic F-127 that were not diluted (Fig. 3). As expected, we observed that increasing the nozzle pressure generally increased the layer width for any given printing speed. In contrast, the output layer width does not significantly change when altering the printing speed for either Pluronic F-127 (Fig. 3a) or collagen bioink (Fig. 3b)

**Figure 3.**
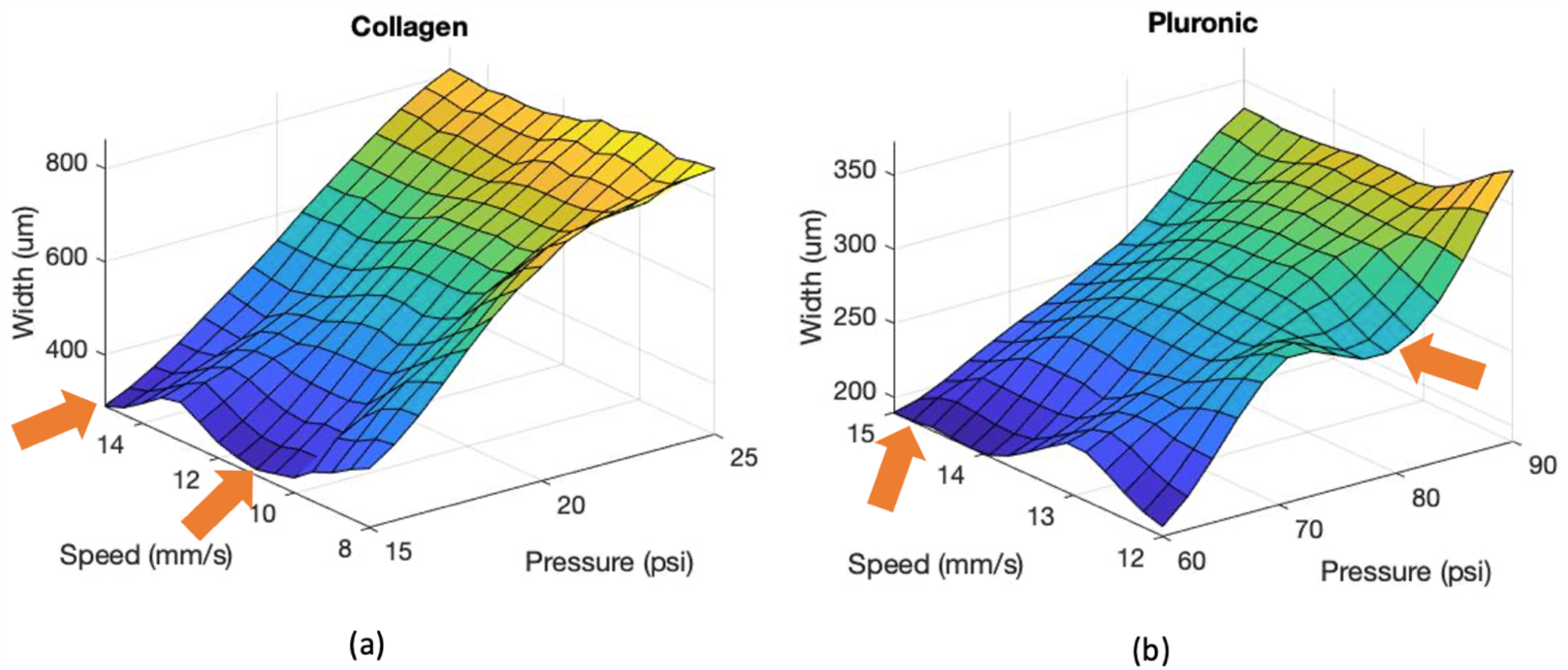
Fuzzy system output for undiluted Pluronic F-127 and collagen bioink. The general shape of the 3D graphs of Collagen (a) and Pluronic (b) shows that increasing the pressure will increase the output layer width. However, at the same pressure level on the 3D graphs of Pluronic and Collagen, the output layer width does not change significantly by expanding the bioprinting speed ratio.

### 3.2 Diluted bioinks are more precise for bioprinting

Next, we simulated the use of each bioink when laden with cells by diluting each bioink to 20% or 40% with αMEM culture media. To overcome the technical difficulty of implementing a fuzzy system with three inputs (pressure, speed, and dilution percentage), we utilized our observation that printing speed for both the collagen bioink and Pluronic F-127 had essentially no effect on layer width (Fig. 4). In particular, an analysis of the undiluted fuzzy output indicates maximum linearity with a printing speed of 15 mm/s for both collagen bioink (Fig. 4a) and Pluronic F-127 (Fig. 4b). As a result, we constructed a fuzzy system using models printed at 15 mm/s and visualized the fuzzy system output for the each bioink (Fig. 5).

**Figure 4.**
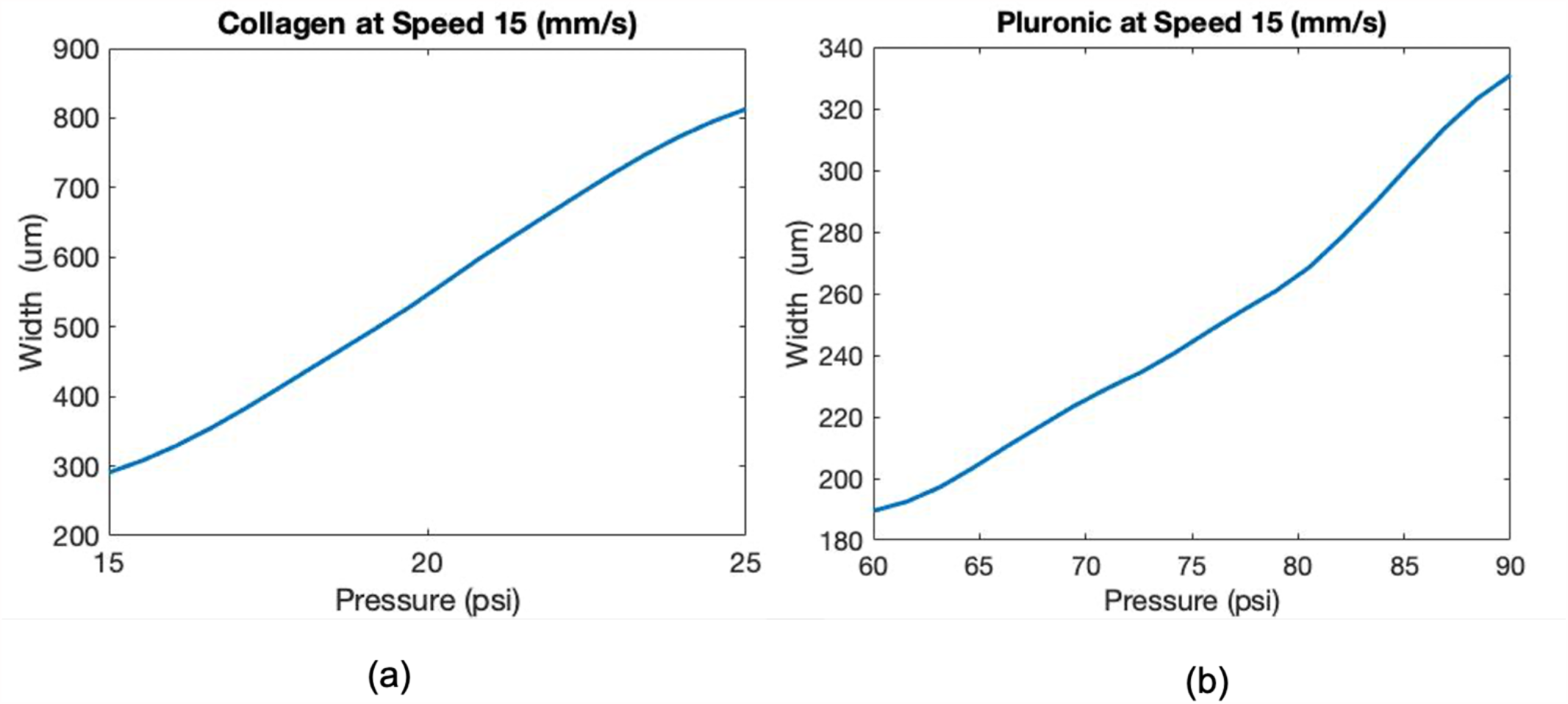
The Pressure linearity at speed 15 mm/s in Collagen and Pluronic. The 2D section of Collagen (a) and Pluronic (b) at speed 15 mm/s show that it has the minimum impact on the output layer width because of the linear relationship between pressure and layer width.

**Figure 5.**
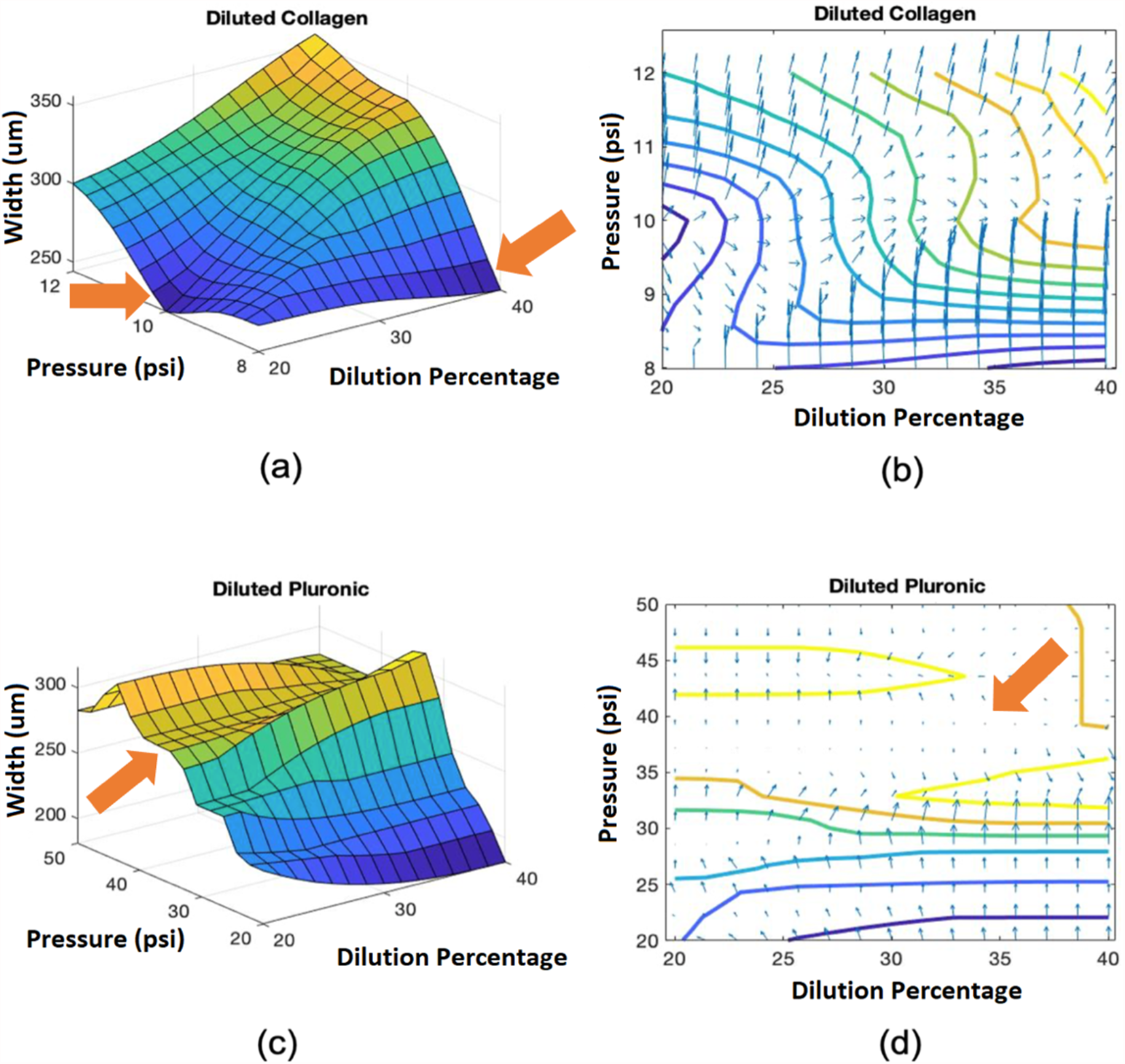
The general shape of the 3D graphs of diluted Collagen and Pluronic with aMEM. The general shape of the 3D graphs of diluted Collagen (a) and diluted Pluronic (c) shows that increasing the pressure will increase the output layer width. (b) and (d) show the gradient based on the explained equation. The smaller arrow inn length is, the bioink has higher printing precision. The flat surface in (c), which is the area between two orange lines (d), indicates a high precision and robust input parameters to print.

In contrast to the undiluted bioinks, the fuzzy output for collagen bioink diluted with αMEM reveals two potential parameter sets that yield a layer width of 200 μm (Fig 5a, illustrated by arrows). The first solution required a high dilution of 40% and a low extrusion pressure of 8 psi. In contrast, the lower dilution of 20% required a slightly higher extrusion pressure of 10 psi to obtain a 200 μm layer width (Fig. 5a). In general, we observed that the layer width increases with increased dilution percentage and extrusion pressure.

In contrast to the undiluted bioink, we observed that the printing precision of diluted collagen bioink was less sensitive to changes in extrusion pressure. In fact, the increase in extrusion pressure from 15 psi to 25 psi increases layer width by 4 fold (37 percent) in the undiluted collagen bioinkas compared to 1.5 fold (9 percent) in the diluted collagen bioink (Fig. 5c). In total, these results indicate that diluted bioinks are more precise for bioprinting.

### 3.3. Bioink Precision Index for novel bionks

Our fuzzy system approach to bioprinting parameter optimization enables us to introduce the Bioink Precision Index (BPI), a new metric for evaluating bioink precision that is defined as the gradient of the Fuzzy 3D surfaces (Fig. 5a,c). The standard calculation for a gradient of a 3D surface is:

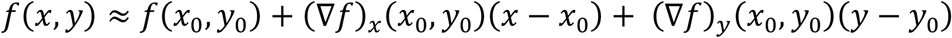

Thus, BPI is calculated from the sum of the squared error of the above equation:

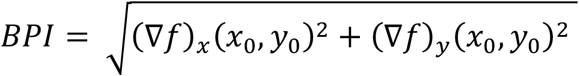

Where *x*_0_ *and y*_0_ are the inputs in our system (speed, pressure, or dilution).

Thus, a smaller numerical value of BPI indicates a more precise parameter set for that bioink. In Fig. 5b and d, BPI is indicated by arrow length at each position. The manufacturer’s suggested printing parameters for collagen bioink (6 mm/s and 15 PSI) results in a BPI of 42.4, but our optimized parameter set (11 m/s and 15 PSI) decreases the BPI to 36, resulting in an approximately 15% increase in precision. In contrast, the BPI for undiluted Pluronic F127 using the manufacturer’s suggested printing parameters (12 mm/s and 80 PSI) is 14.2, much greater than the BPI of 4.5 obtained using our optimized parameter set (15 mm/s 60 psi). Finally, we note that the optimized BPI for collagen is significantly higher than the optimized BPI for pluronic, indicating that collagen is a more challenging bioink to use for high precision bioprinting.

### 3.4. Fuzzy systems identify robust and precise printing parameters sets

A potentially useful feature of the fuzzy system approach is the identification of printing parameter sets that are insensitive to the small perturbations routinely encountered during bioprinting, such as inconsistent experimental dilutions or fluctuations in extrusion pressure during printing. One such parameter set can be visualized by the flat surface in Fig. 5c (arrow), which results in the same 250 μm layer width for any dilution between 40% and 50% and extrusion pressure between 20 psi and 40 psi.

To illustrate these areas of Fig. 3a, we separately analyzed sections illustrating the speed-width relationship for a given a pressure (Fig. 6a) and pressure-width relationship for a given speed (Fig. 6b). Since BPI can be reduced to one dimension (one input-output system) or higher dimensions, we calculated the BPI for one dimension model (pressure-width). The figures are color-coded with green, yellow, or red, where the green zone has the most precision and least sensitivity to fluctuations, resulting in the most precise outputs (BPI < 1). In contrast, the yellow (1 < BPI < 20) and red (BPI > 20) zones are progressively more challenging to achieve a high level of precision over time. Similar regions have been identified for undiluted Pluronic F-127 (Fig. 6c,d), diluted collagen bioink (Fig. 6e, f), and diluted Pluronic F-127 (Fig. 6g, h).

**Figure 6.**
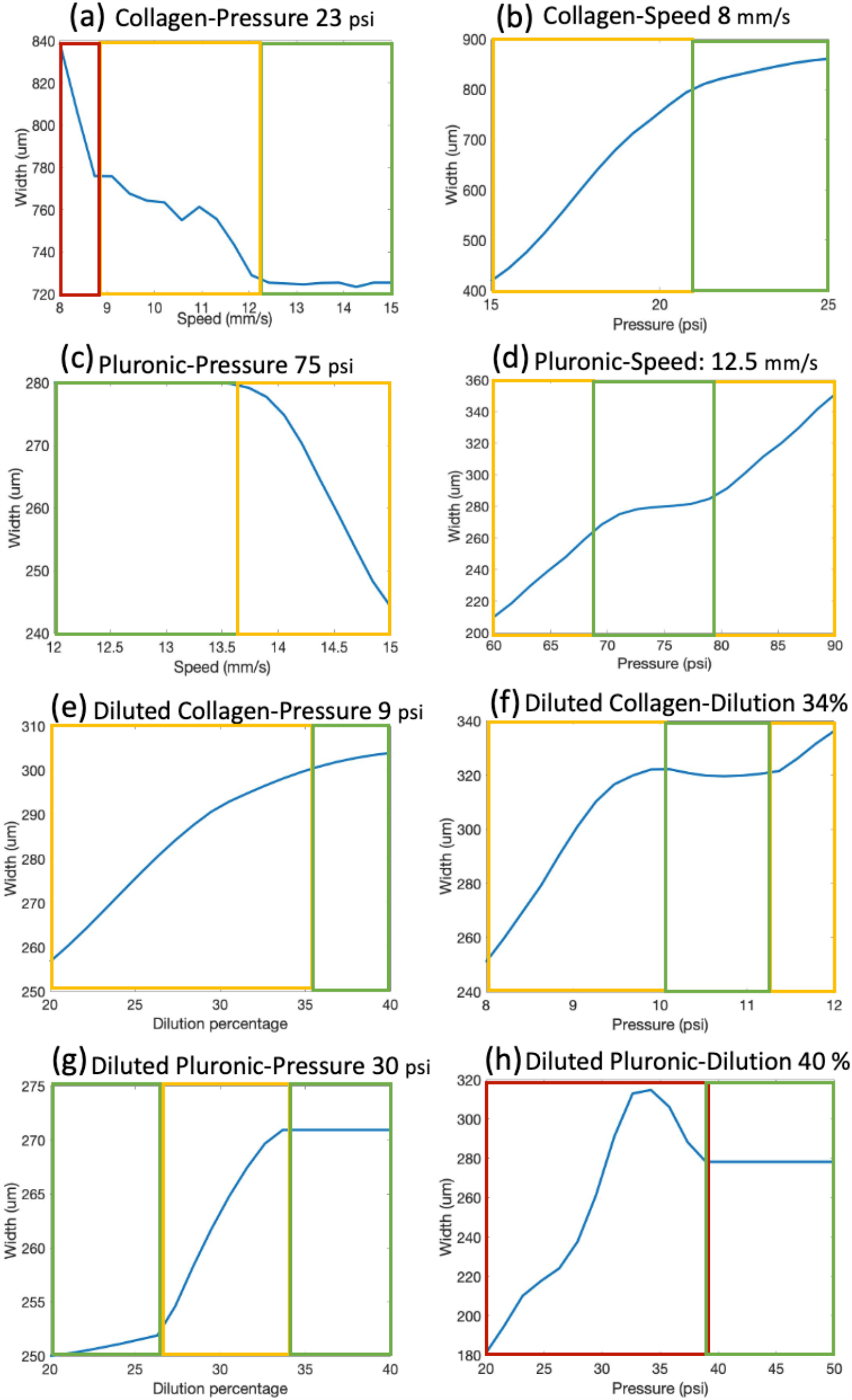
The 2D slices of Collagen, Pluronic, diluted Collagen, and Pluronic with aMEM. This figure indicates the 2D slices in x and y direction at specified value written as the label for the Collagen (a and b), Pluronic (c and d), diluted Collagen with aMEM (e and f), and diluted Pluronic with aMEM (g and h).

## 4. Discussion

Currently, there is no method to evaluate bioink printability and precision. Here, we have shown that fuzzy systems can be used to identify optimized parameter sets that can be used to improve the precision of existing bioinks. In this study, we found that fuzzy optimization improved precision in collagen bioink by 15%, as compared to the manufacturer’s recommended printing parameters. Furthermore, we have introduced a new standardized metric (BPI) that can be easily generated and is useful for comparing precision between bioinks. Here, we used BPI to illustrate that collagen bioink is more challenging for precision printing than pluronic bioink and that diluted bioinks are more precise than non-diluted bioinks.

The fuzzy system developed here identified one or multiple sets of optimized parameters for each bioink. We note that the target layer width for our model was 200 μm. As illustrated in Fig. 3a (arrow), collagen bioink would meet this requirement at the low pressure of 15 psi and speed of 10 or 15 mm/s. In contrast, printing undiluted Pluronic F-127 at 200 μm layer width is possible with two different parameter sets, as illustrated in Fig. 3b (arrows). Here, we observed that it is possible to meet this requirement with either high pressure (80 psi) and low speed (12 mm/s) or low pressure (60 psi) and high speed (15 mm/s). Generally low pressure is preferred due to increased cell viability [22], so this observation could significantly improve printing time and experimental outputs when using Pluronic F-127. Investigators are likely to make similar observations if this approach was applied during novel bioink development.

Previous attempts to improve bioprinting precision have focused on developing novel bioprinting techniques, such as the miniaturized progressive cavity pump method to replace the extrusion-based method [23]. However, utilizing this soft computing technique increases bioprinting precision in addition to any improvements to the method itself. Moreover, the fuzzy surface is useful for understanding tradeoffs between precision and other constraints (e.g. cells sensitive to high pressure). Nonetheless, our model consisted of three inputs (pressure, speed, and dilution) and one output (layer width) that are directly related to the extrusion bioprinting method. However, future work may extent this approach to other parameters that affect bioprinting or even experimental outcomes, such as cell viability or biocompatibility. Finally, we note that BPI is independent from input dimensions/units and will remain a useful metric for comparison between bioinks as bioprinting technology evolves.

## 5. Conclusion

Obtaining high precision in bioprinting is a necessary step towards mass production of bioprinted constructs for use in research and medicine. Here, we have demonstrated that a fuzzy system can be used to approximate layer width given a set of bioprinting parameters, including printing speed, extrusion pressure, and media dilution percentage, as well as determine bioprinting parameter sets that maximize precision. Furthermore, we have defined the Bioink Precision Index (BPI) that can be used to quickly compare the ease of reproducibility across the wide variety of bioinks currently available.

